# Cytogenetic evidence supports *Avena insularis* being closely related to hexaploid oats

**DOI:** 10.1101/2021.08.25.457631

**Authors:** Araceli Fominaya, Yolanda Loarce, Juan M. González, Esther Ferrer

## Abstract

Cytogenetic observations, phylogenetic studies and genome analysis using high-density genetic markers have suggested a tetraploid *Avena* species carrying the C and D genomes (formerly C and A) to be the donor of all hexaploid oats (AACCDD). However, controversy surrounds which of the three extant CCDD tetraploid species - *A. insularis, A. maroccana* and *A. murphyi* - is most closely related to hexaploid oats. The present work describes a comparative karyotype analysis of these three CCDD tetraploid species and two hexaploid species, *A. sativa* and *A. byzantina*. This involved the use of FISH with six simple sequence repeats (SSRs) with the motifs CT, AAC, AAG, ACG, ATC and ACT, two repeated ribosomal sequences, and C genome-specific repetitive DNA. The hybridization pattern of *A. insularis* with oligonucleotide (AC)_10_ was also determined and compared with those previously published for *A. sativa* and *A. byzantina*. Significant differences in the 5S sites and SSR hybridization patterns of *A. murphyi* compared to the other CCDD species rule out its being directly involved in the origin of the hexaploids. In contrast, the repetitive and SSR hybridization patterns shown by the D genome chromosomes, and by most of the C genome chromosomes of *A. maroccana* and *A. insularis*, can be equated with the corresponding chromosomes of the hexaploids. Several chromosome hybridization signals seen for *A. insularis*, but not for *A. maroccana*, were shared with the hexaploid oats species, especially with *A. byzantina*. These diagnostic signals add weight to the idea that the extant *A. insularis*, or a direct ancestor of it, is the most closely related progenitor of hexaploid oats. The similarity of the chromosome hybridization patterns of the hexaploids and CCDD tetraploids was taken as being indicative of homology. A common chromosome nomenclature for CCDD species based on that of the hexaploid species is proposed.

## Introduction

Hexaploid *Avena* species (2n = 6x= 42), including cultivated *A. sativa* and *A. byzantina*, have three genomes of seven chromosomes each. Studies on the genome constitution of these species [1, 2] have used a formula based on the putative origin of these genomes. Accordingly, the hexaploids arose from an original hybridization between a tetraploid species with the genomes A and C, and a diploid species with a D genome. These studies were mainly based on karyotype analyses involving conventional staining [3], C-banding [4–13] and fluorescent *in situ* hybridization (FISH) with repetitive DNA probes [14–20]. However, no definite correspondence among chromosomes from diploid, tetraploid and hexaploid species has yet been confirmed. Analyses of chromosome pairing in interspecific hybrids showed a departure from the expected bivalent number, although close homology among chromosomes of the different ploidy number species was maintained (revised in [21]. The detection of multivalents during meiosis in interspecific hybrids indicates that the extant *Avena* species differ in their chromosome structure, mainly a consequence of translocations and inversions that occurred during their evolutionary history. Genome *in situ* hybridization (GISH) has shown that small C genome segments are translocated onto the A or D genome chromosomes and *vice versa* [22–25]. FISH with genome-specific repetitive sequences has identified chromosomes involved in translocations [14, 15, 26] in both tetraploid and hexaploid species. Changes in these chromosome structures have also been deduced through comparisons of genetic maps of diploid, tetraploid and hexaploid species [27–29]. As a result of these profuse rearrangements, none of the extant diploid species has been unequivocally identified as the donor of the A, C and D genome.

The results of the above studies have led to the general agreement that no extant diploid species possesses the D genome as it exists in the tetraploid and hexaploid species. However, growing evidence supports the idea that tetraploid AACC species contain the genome designated as D in hexaploid oats. Many studies have suggested that the D genome may have originated from an A genome species given the close relationship between them [30–32]. However, FISH analysis using a specific repetitive sequence of the A genome, discriminated among the chromosomes of the A and D genomes in tetraploids and hexaploids [16]. Moreover, chromosome morphology, and the distribution of chromosome markers (such as ribosomal loci and translocated segments), showed a close resemblance between certain chromosome pairs of the A genome of tetraploids and the D genome of hexaploids. Based on these observations, Fominaya et al. [33] suggested that tetraploids likely carried a D genome instead of an A genome More recently, phylogenetic studies of the genus comparing nucleotide sequences from chloroplasts and nuclear gene sequences [34–36], as well as genome wide analysis [37, 38] have supported this idea. Moreover, a comparison of the chromosome distribution of repetitive sequences among tetraploid and hexaploid species has strongly suggested that the D genome is present in these tetraploids [39].

Further discrepancies exist over whether one of the three known CCDD tetraploids - *A. insularis* Ladiz., *A.maroccana* Gdgr (synonym *A. magna* Murphyi et Terr) and *A. murphyi* Ladiz. - or an extinct tetraploid species, might have been the immediate tetraploid ancestor of hexaploid oats. *A. insularis* has been proposed as this ancestor based on chromosome morphological similarities [9] and hybrid pairing data [21], with additional evidence provided by genetic diversity studies using SNPs [27] and high-density genetic markers revealed by genotyping-by-sequencing (GBS) [38]. However, phylogenetic relationships in the genus *Avena* based on the ITS of 45S rDNA and the nuclear *Pgk1* gene (widely used to reveal the evolutionary story of other grass species) suggest a closer relationship of the hexaploids with *A. maroccana* and *A. murphyi* than with *A. insularis* [40, 41].

Microsatellites or simple sequence repeats (SSRs) are tandemly repeated sequences in which the repeated unit covers 1-6 bp. Importantly, microsatellites are found throughout the genome, with differences in the number of repeated units at each location leading to high levels of polymorphism. Short microsatellite tandems are commonly used as genetic markers for the study of many species. In *Avena* they have been used to infer genetic relationships within species, among closely related species [42] and in the construction of linkage maps [43–45]. Since microsatellites can also organize themselves into large arrays containing thousands of units, they can also be used as physical markers. Indeed, since the work of Cuadrado and Schwarzacher [46], oligonucleotides containing a few copies of the repeated unit have been much used as probes in FISH experiments for identifying and karyotyping plant chromosomes. This methodology overcomes the need for cloning repetitive sequences and increases the number of available chromosome markers.

The distribution patterns observed for SSRs [47–48] are not so different from those of the known satellite DNA families. These satellites are often common to related species, whereas some differ considerably between species. In general, satellite sequences are more similar among closely related species than among distant species, but the content and diversity of tandem repeated DNA can differ even in closely related species [49–50]. Thus, the study of distribution patterns can detect chromosome variation in terms of the abundance and distribution of a repeat among close related species - which is of great interest when trying to determine the genome constitution of polyploid species and the origin of their constituent genomes.

Comparative SSR-FISH karyotyping has been performed with many different grass species, including wheat [47, 51], barley [52, 53] and *Avena* species [39, 54–58,]. These studies differed in the number of species and SSRs analyzed, although all of them confirmed the validity of the strategy for karyotyping *Avena* chromosomes and detecting modifications in the chromosome structure among diploid and polyploid species. However, no diagnostic chromosome markers that could help trace the origin of the different genomes in polyploid species were clearly revealed.

With respect to the relatedness of CCDD tetraploid species and hexaploids, FISH mapping of the AC microsatellite sequence has indicated proximity between the chromosomes of *A. maroccana* and *A. sativa*, with more distant hybridization patterns observed between *A. murphyi* and *A. sativa* [54] (*A. insularis* was not included in the analysis). In their study of the distribution patterns of three SSRs, TTC, AAC and CAC, Yan et al. [39] found no FISH-derived tetraploid karyotype closer than any other to the hexaploids. In contrast, Luo et al. [57] suggested that *A. insularis* had the most similar ACT pattern to the hexaploids, although no precise correspondence between the D chromosomes of the two polyploid species was seen.

In the present study, the FISH distribution patterns of six SSRs, namely CT, AAC, AAG, ACG, ACT and ATC, were used in FISH experiments with the three CCDD species *A. insularis*, *A. maroccana* and *A. murphyi*, and the two hexaploid species *A. byzantina* C. Koch and *A. sativa* L. In addition, AC was analyzed in *A. insularis* and its distribution pattern was compared with the other CCDD tetraploids and hexaploid species previously studied [54]. The results suggest *A. insularis* to be the CCDD species most closely related to the *Avena* hexaploids.

Taking into account the possible similarities among the chromosome of tetraploid and hexaploid species, it is here tentatively proposed that homologous relationships exist between specific tetraploid and hexaploid chromosomes. A common terminology for CCDD tetraploid species chromosomes is proposed on the basis of that used by Sanz et al. [18] for hexaploids.

## Materials and Methods

### Plant Materials

Table 1 shows the *Avena* species used in the present study. These included three tetraploid forms with a CCDD genome constitution, and two hexaploid species with an AACCDD constitution. These species were kindly provided by different germplasm resource centres.

**Table 1.**
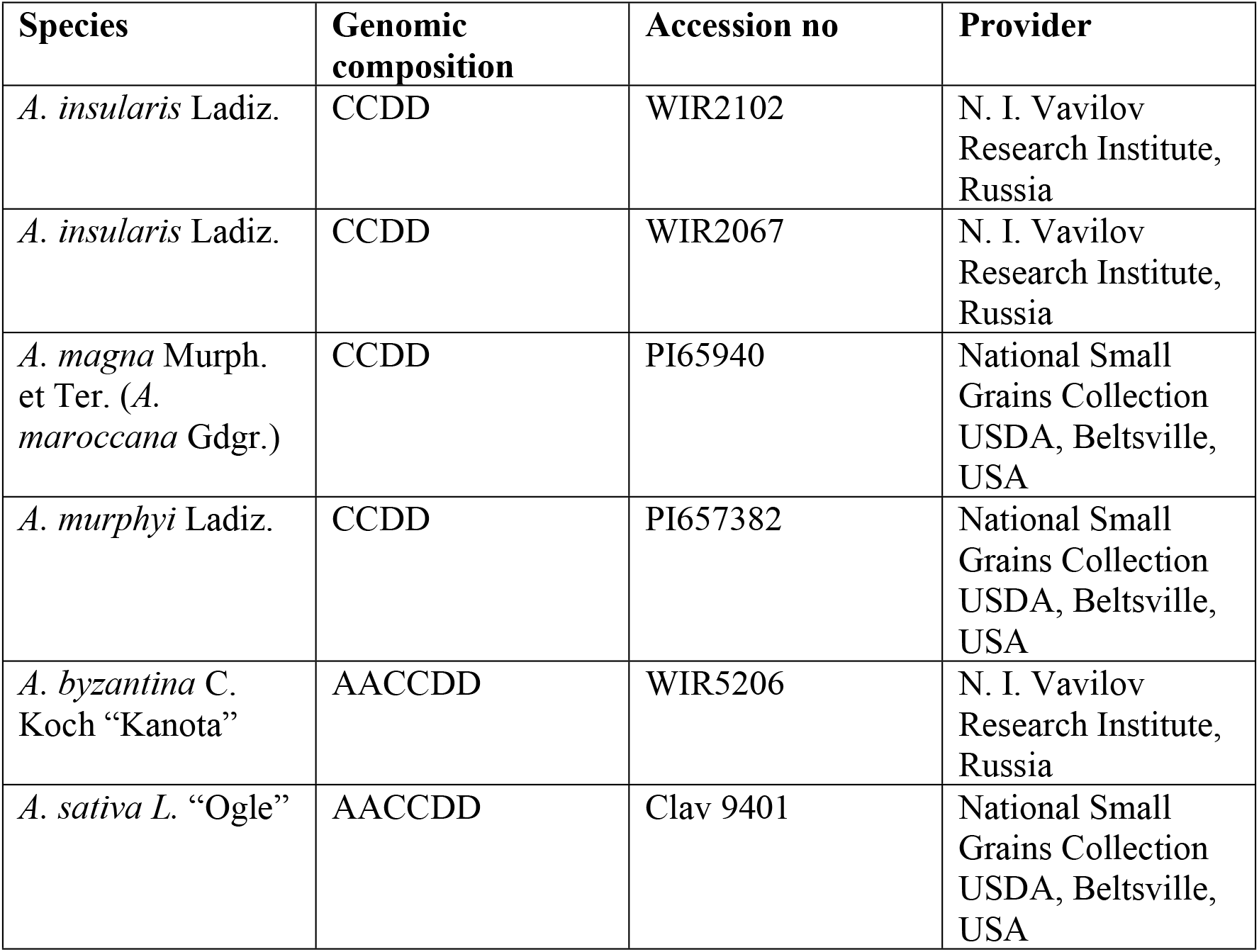
Plant material used in the present study.

#### Mitotic metaphase preparations

Root tips were obtained from seedlings, and mitotic metaphases prepared as previously described [14, 54].

#### Probes and FISH

Ten synthetic oligonucleotides were initially used. Two of these contained di-nucleotide motifs, and eight contained tri-nucleotide motifs (S1 Table). All were labelled with biotin at both ends. Four repetitive probes were used for chromosome identification: (1) pAs120A, specific for the oat A genome, containing an insert of 114 bp isolated from *A. strigosa* [16]; (2) pAm1, specific for the oat C genome, containing an insert of 464 bp isolated from *A. murphyi* [59]; a biotin-labelled oligonucleotide containing 51-mers derived from the Am1 sequence was alternatively used [55]; (3) p45S, a ribosomal probe derived from *A. strigosa* [18] and (4) pTa794, containing an insert of 5S rDNA isolated from *Triticum aestivum* [60]. All probes were labelled with digoxigenin-11-dUTP or biotin-16-dUTP.

Prior to FISH, chromosome preparations were treated with 4% (w/v) paraformaldehyde and dehydrated in an ethanol series followed by air drying. SSR-FISH was performed as described by Fominaya et al. [54]. The hybridization mixture (30 μL) contained 50% (v/v) formamide, 2xSSC, 10% (w/v) SDS, 10% dextran sulphate, 50 μg/mL of *Escherichia coli* DNA, and 2.6 pmol of the SSR. Hybridization was performed at 37°C overnight. Post-hybridization washing was performed in 4xSSC/0.2% Tween-20 for 10 min at room temperature (RT). The detection of labelling and FISH sequential hybridization with repetitive probes were performed as described previously [16, 54].

Images were obtained using a Zeiss Axiophot epifluorescence microscope. Images captured from each filter were recorded separately using a cooled CCD camera (Nikon DS) and the resulting digital images processed using Adobe Photoshop. For each combination, at least three slides were studied, with 5-10 metaphase cells analyzed per slide.

## Results

### (AC)_n_-FISH signal pattern in *A. insularis*

The karyotype of *A. insularis* was described by [9] using C-banding. FISH with ribosomal probes to detect the 45S and 5S loci, plus repeated sequences specific to the A and C genomes (As120a and Am1, respectively) was carried out by Fominaya et al. [33] although a complete assignment of chromosomes was not provided in that work. Thus, for a complete description of the *A. insular*i*s* standard karyotype, the chromosome numbering used was that proposed by Jellen and Ladizinsky [9] which was based on relative chromosome length and arm ratios, together with the information provided here by the location of ribosomal probes and hybridization patterns with pAm1.

No polymorphic hybridization signals were detected between the two *A. insularis* accessions with any of the probes or oligonucleotides used. FISH with pAs120a failed to reveal hybridization on any chromosome of *A. insularis* (data not shown), whereas FISH with pAm1 detected a dispersed pattern of hybridization on 14 chromosomes that corresponded to the C genome (Fig 1a and d). Several C chromosomes, namely M1, M2, and SM3, showed no hybridization on the terminal regions of their long arms, indicating translocations from D chromatin. Other putative interstitial translocations in the long arms were identified in SM1 and SM2. A large pAm1 hybridization signal was located at the terminal region of the long arm of chromosome SM1, rendering this chromosome easily identifiable. In contrast, the D genome chromosomes showed hybridization with pAm1 for only four chromosome pairs. SM5, SM6, SAT1 and SAT2 showed terminal hybridization signals of different intensity with this specific C genome probe, all diagnostic of intergenomic C/D translocations (Fig 1a, 1d and S1 Fig). Among these, the most intense hybridization signal was that seen for SAT2, and the least intense for SAT1. Both chromosomes had 45S loci, and SAT1 also carried two 5S loci on its long arm (Figs 1b and d). Two C genome chromosomes, M1 and M2, also had 5S loci on their long arms.

**Fig 1.**
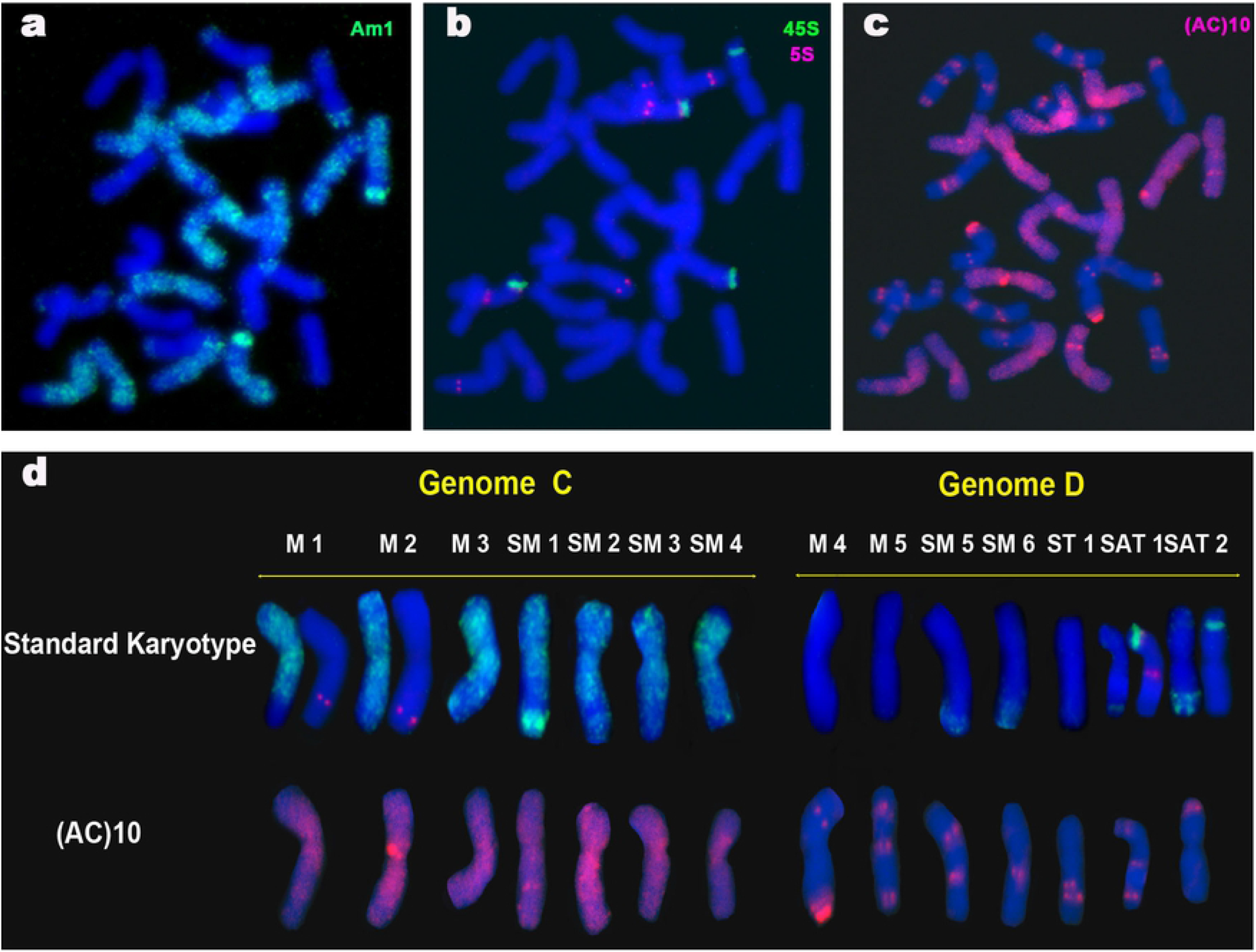
FISH of a mitotic metaphase cell of *A. insularis* showing the distribution of repeated sequences Am1, ITS 45S, 5S and AC. (a) Am1 (green). (b) 45S (green) and 5S (red). (c) SSR AC (red). (d) *A. insularis* karyotype showing a single chromosome of each homologous pair. The standard karyotype is based on the distribution patterns of Am1 (green), ITS 45S (green) and 5S (red). Chromosome nomenclature is based on that of Jellen and Ladizinsky [9].

Hybridization with oligonucleotide (AC)_10_ gave very different patterns for the C and D genome chromosomes (Figs 1c and d). All C genome chromosomes showed conspicuous AC signals in the pericentromeric regions, covering areas of variable length on both arms. The strongest were on chromosomes M2 and M3. However, the hybridization pattern obtained with (AC)_10_ added almost no new information to the standard karyotype in terms of differentiating each individual C genome chromosome pair. Only a conspicuous signal observed interstitially on the long arm of chromosome SM1 allowed this chromosome to be undoubtedly identified. In contrast, and similar to that described for other tetraploid and hexaploid *Avena* species [54]), the D genome chromosomes that hybridized with (AC)_10_ showed discrete signals of different intensity. Indeed, (AC)_10_ unequivocally distinguished the seven D genome chromosomes of this species. A combination of either subtelomeric, interstitial and pericentromeric signals on one or both arms of each chromosomes were shown by each chromosome pair (Fig 1d).

### SSR-FISH signal patterns in tetraploid species

The three CCDD tetraploid species were analyzed by FISH using nine oligonucleotide SSR probes, although only six of them produced discrete hybridization signals (S1 table). Hybridization patterns of these six SSRs are shown in Fig 2.

**Fig 2.**
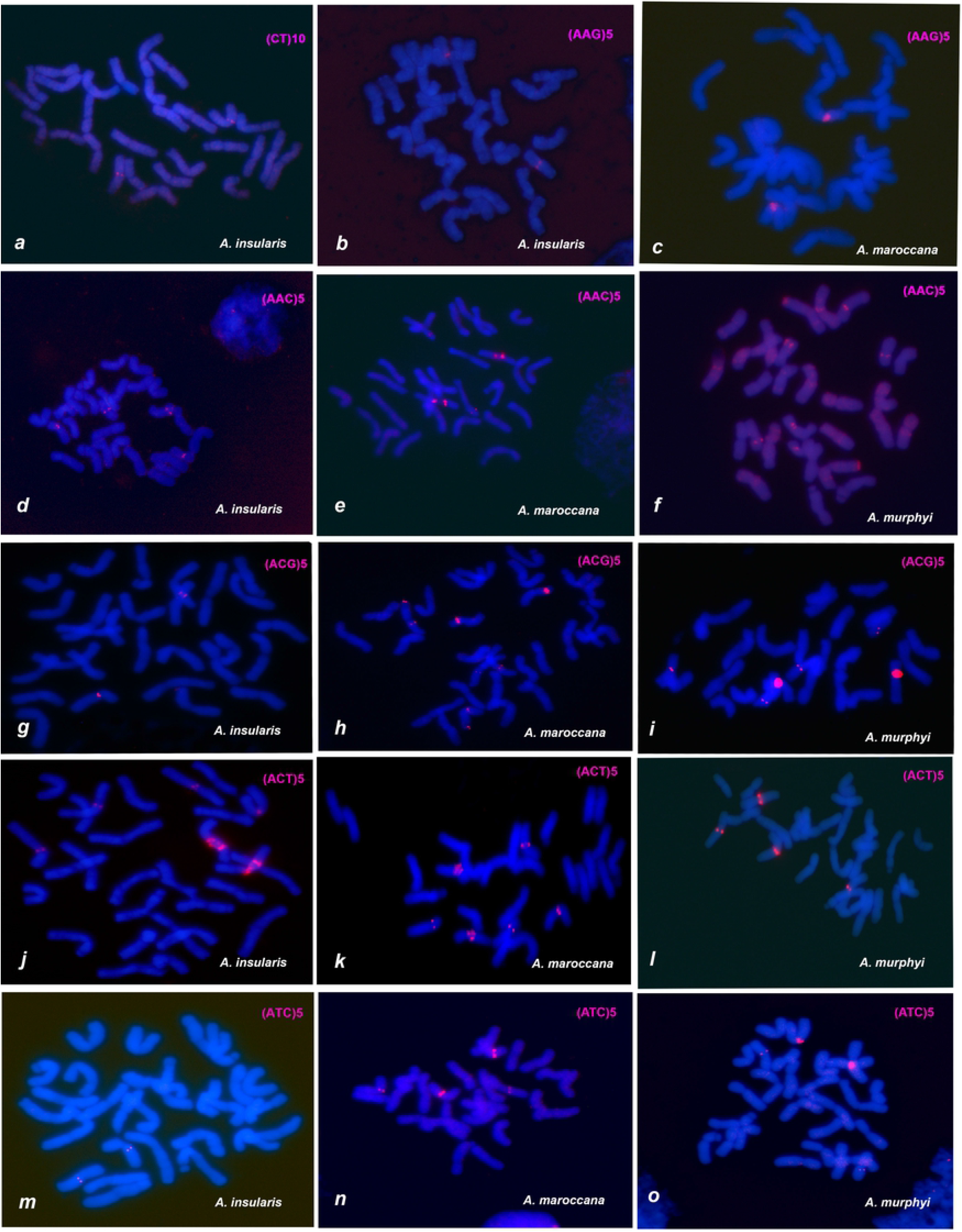
FISH of mitotic metaphases of CCDD tetraploid species, *A. insularis, A. maroccana* and *A. murphyi*, showing the distribution of SSR hybridization signals in red. (a) CT. (b and c) AAG. (d-f) AAC. (g-i) ACG. (j-l) ACT. (m-o) ATC.

FISH analysis using each single SSR probe was followed by re-hybridizations of the same metaphases with probes containing 5S, 45S and Am1 repetitive sequences to facilitate chromosome identification (S2 and S3 Figs). Karyotypes of each species showing diagnostic hybridization signals were obtained (Fig 3).

**Fig 3.**
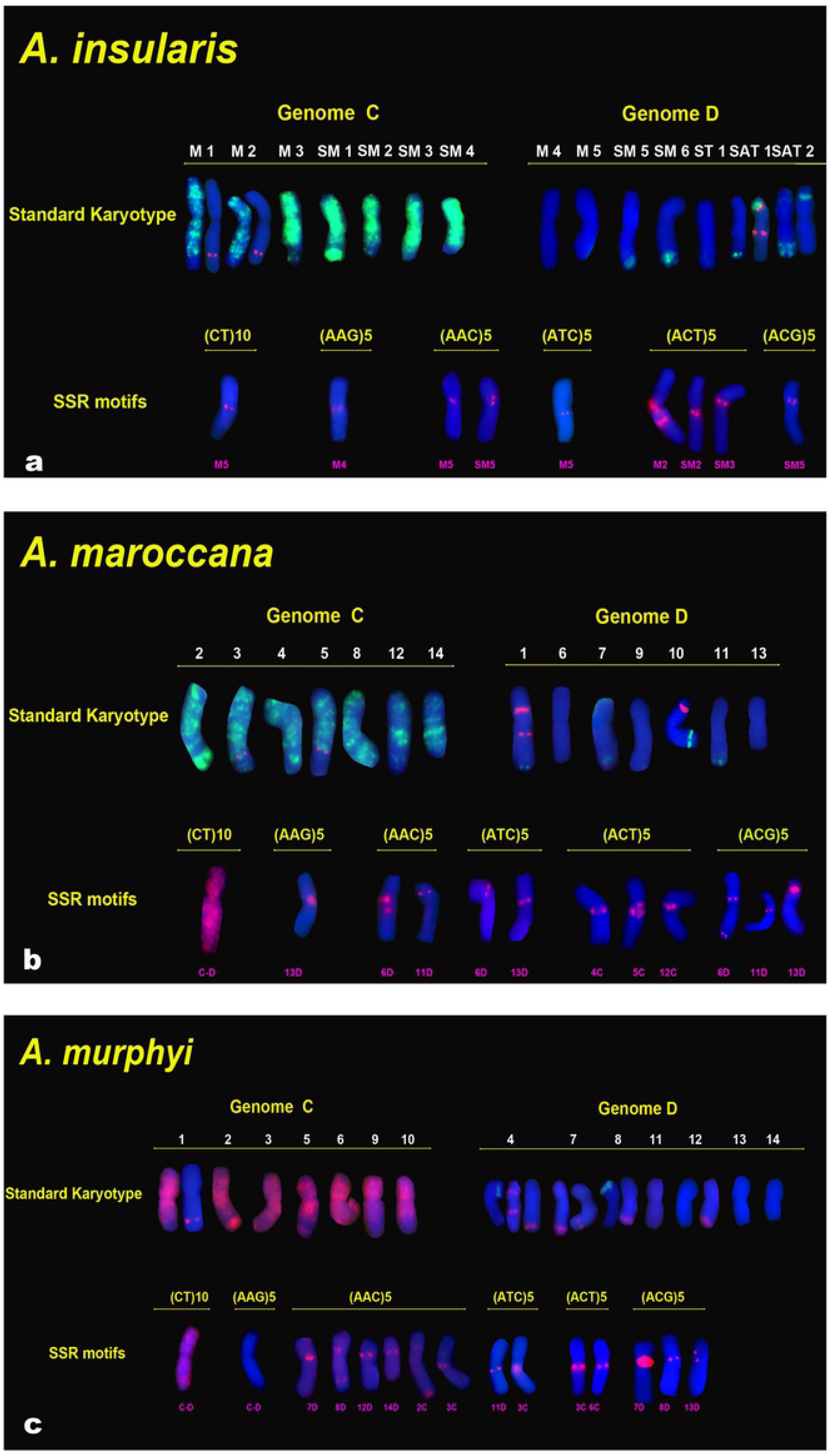
Karyotypes of CCDD tetraploid species showing a single chromosome of each homologous pair from the metaphases of cells in Fig 3. Standard karyotypes are based on the FISH distribution patterns of Am1, ITS and 5S. Chromosome nomenclature is based on that of Jellen and Ladizinsky [9] for *A. insularis* and Fominaya et al. [5, 54] for *A. maroccana* and *A. murphyi*. (a) *A. insularis* standard karyotype: Am1(green), 45S (green, Sat1 and Sat2) and 5S (red, 1C, 2C and Sat1). (b) *A. maroccana* standard karyotype: Am1 (green), 45S (red, 1D and 10D) and 5S (red, 3C, 5C and 1D). (c) *A. murphyi* standard karyotype: Am1 (red), 45S (green, 4D and 8D) and 5S (red, 1C, 4D and 7D). Panels of SSR motifs show chromosomes with specific SSR signals in red.

Chromosomes of *A. insularis* were numbered following the nomenclature described above (Fig 3a). Chromosomes of *A. maroccana* (Fig 3b) and *A. murphyi* (Fig 3c) were numbered 1-14 according to the nomenclature of Fominaya et al. [5, 54]. These last two nomenclatures were based on chromosome arm ratios, relative lengths, and the FISH patterns for the three repetitive probes. It is worth mentioning that the hybridization of *A. maroccana* with pAm1 revealed four terminal C/D intergenomic translocations on chromosomes 1D, 7D, 10D and 11D (Figs 3b, S1 and S2 Figs) similar to that seen in *A. insularis*. In contrast, *A. murphyi* showed two clear C/D translocations on chromosomes 7D and 8D, and two minor ones on chromosomes 4D and 12D that were visible after increasing the CCD exposure time (Figs 3c, S1 and S3 Figs.). 45S loci for both species were found in locations previously described [26] (Figs 3b and 3c). Loci for 5S were identified in *A. maroccana* (Figs 3b, S3 Fig) similar to that described here for *A. insularis* - namely, two loci on the 3C and 5C chromosomes, and a double signal on chromosome 1D. In *A. murphyi*, four 5S loci were observed but with a distribution slightly different to that previously described [26] (Fig. 3c): a double signal was present on the satellited chromosome 4D as in the other two species, but single signals only were seen on chromosome 1C and chromosome 7D. Chromosome 7D had a C/D translocation, and the 5S signal lies close to it.

In contrast to the hybridization pattern shown by (AC)_10_ in *A. insularis* (Fig 1), and the patterns previously described for the other two CCDD species [54], none of the six SSRs analyzed produced signals on all 14 chromosome pairs of either CCDD species (Figs 2 and 3). The SSRs studied produced signals on a few chromosome pairs within each species and most produced signals exclusively on chromosomes of one genome. D genome chromosomes were the best represented in the hybridization patterns, with signals located in the centromeric and pericentromeric regions. Oligonucleotide (CT)_10_ returned a centromeric signal on chromosome pair M5 of the D genome of *A. insularis* (Figs 2a), whereas it was dispersed throughout the chromosomes of *A. maroccana* and *A. murphyi* with no discrete signals at all (S2 Fig). The absence of a discrete signal was also observed in *A. murphyi* for AAG (S2 Fig), although a centromeric signal was seen on the M4 chromosome pair of *A. insularis* (Fig 2b) as well as on chromosome 13D of *A. maroccana* (Figs 2c). Oligonucleotide (AAC)_5_ revealed significant differences between *A. murphyi* and both *A. insularis* and *A. maroccana*. Only two pairs of D chromosomes showed signals with this oligo, both in *A. insularis*, (M5 and SM5) (Fig d) and in *A. maroccana* (6D and 11D) (Fig 2e). In *A. murphyi*, however, four D genome chromosome pairs (7D, 8D, 12D, and 14D) and two C genome chromosome pairs (2C and 3C) showed pericentromeric or telomeric signals (Figs 2f). Differences in signal intensity for this SSR between *A. insularis* and *A. maroccana* were notable. A double strong pericentromeric AAC signal was evident on chromosome 6D of *A. maroccana* (Fig 2c) while only single weaker signals were observed on M5 and SM5 of *A. insularis* (Fig 2d). Distribution of ATC showed differences both in terms of signal intensity and the presence/absence of discrete signals (Figs 2m and 2n). Oligonucleotide (ATC)_5_ produced a very faint centromeric signal on the D genome chromosome M5 of *A. insularis* (Fig 2m), and more intense on 6D and 13D of *A. maroccana* (Fig 2n). In *A. murphyi*, and similar to that described for oligonucleotide (AAC)_5_, ATC signals were observed on chromosomes of the two genomes, 3C and 11D (Fig 2o). Significant differences among *A. insularis* and the other two species were observed for ACG signals. In the three species, signals appeared exclusively on D genome chromosomes, but in *A. insularis* only the chromosome pair SM5 showed pericentromeric signals (Fig 2g), while three chromosome pairs of *A. maroccana* (6D, 11D and 13D) (Fig h) and *A. murphyi* (7D, 8D and 13D) showed them (Fig 2i). Unlike the distribution patterns of the other SSRs studied, oligonucleotide (ACT)_5_ produced pericentromeric hybridization signals of variable intensity only on chromosomes of the C genome in the three tetraploids. *A. insularis* and *A. maroccana* showed similar signals on three chromosome pairs (M2, SM2 and SM3) in *A. insularis* (Fig 2j) and in *A. maroccana* (4C, 5C and 12C) (Fig 2k). However, only two chromosome pairs (3C and 6C) showed ACT signals in *A. murphyi* (Fig 2l).

The pericentromeric regions of the chromosomes of the CCDD species seemed to be enriched in different SSR sequences. Co-located signals were evident after FISH with different oligonucleotides (Fig 3). For example, in *A. insularis*, CT, AAC and ATC co-localized on chromosome M5, or in *A. maroccana*, AAG, ATC, and ACG co-localized to 13D (Figs 3a, 3b). However, within a species, pairs of SSRs did not necessarily hybridize to the same chromosomes, indicating a specific combination of SSRs to be present at each centromeric or pericentromeric location. For instance, not all chromosomes of *A. maroccana* with signals for ACG showed signals for ATC (Figs 2h 2n and 3b), and in *A. murphyi* AAC and ATC signals did not always coincide at the same chromosome locations (Figs 2f, 2o and 3c). However, certain SSR combinations were common to the chromosomes of different species, especially those of *A. insularis* and *A. maroccana*. For instance, M5 of *A. insularis* and 6D of *A. maroccana* shared the signal combination for AAC and ACG; the same was true for SM5 of *A. insularis* and 11 D of *A. maroccana* (Figs 3a and 3b).

Despite the small number of chromosomes with identifiable FISH signals produced by hybridization with each SSR, interspecies differences for each single SSR were observed in terms of the number of chromosomes with signals, and the signal intensity (Fig 3). On the whole, *A. insularis* showed fewer and more weak hybridization signals than the other two species for most of the SSRs, while *A. murphyi* showed the most different hybridization patterns with respect to the other two CCDD tetraploids. Although *A. insularis* and *A. maroccana* showed few discordant hybridization patterns, clear differences between them were evident (Fig 3a and 3b).

### SSR-FISH signal patterns in hexaploid species

The hexaploid species *A. byzantina* and *A. sativa* were karyotyped with the same six oligonucleotides previously employed with the tetraploid species. SSR hybridization patterns are shown for the oligonucleotides (CT)_10_, (AAC)_5_, (AAG)_5_, (ACG)_5_, (ACT)_5_ and (ATC)_5_ (Fig 4).

**Fig 4.**
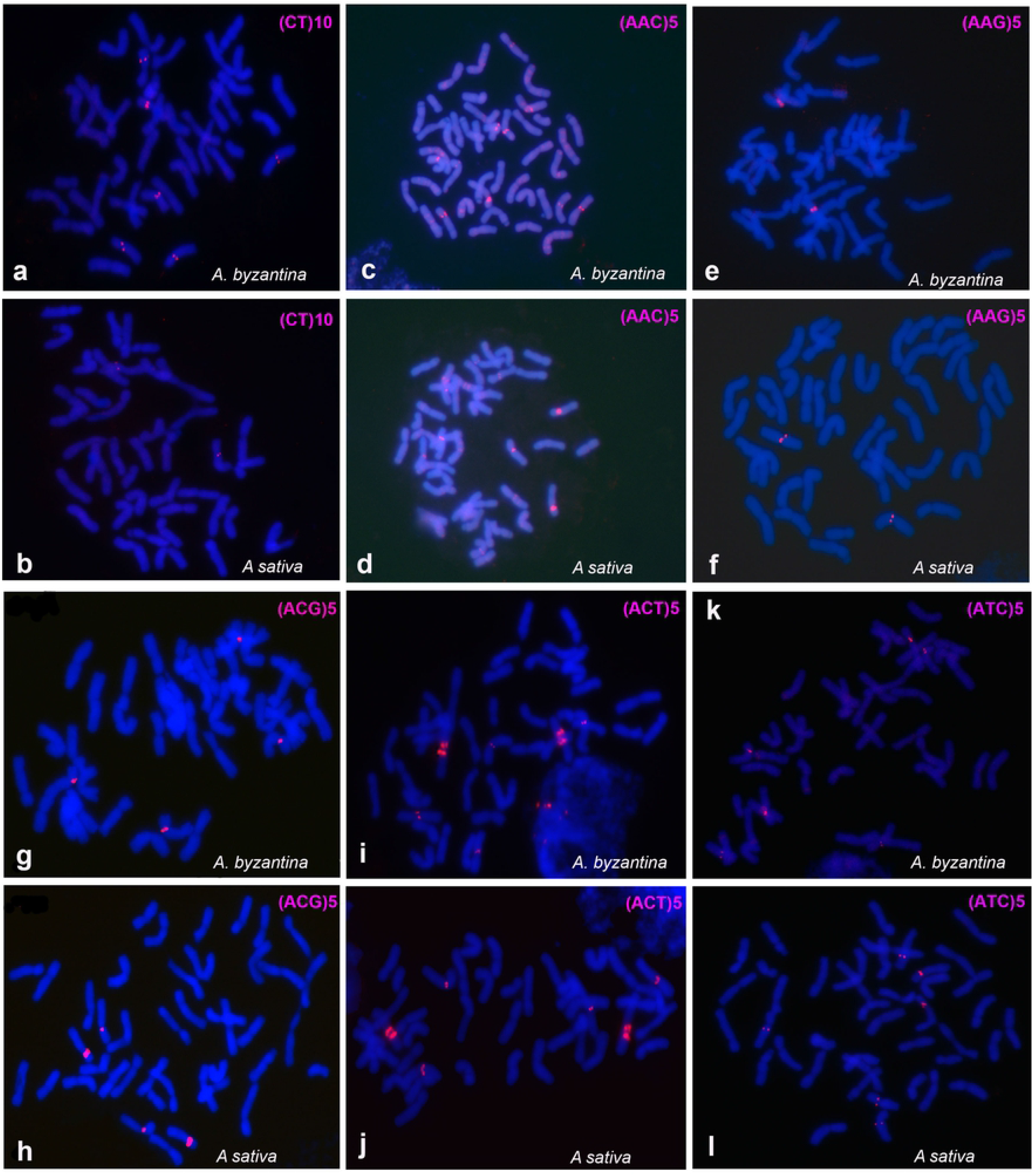
FISH of mitotic metaphases of hexaploid species, *A. byzantina* y *A. sativa*, showing the distribution of SSR hybridization signals in red. (a and b) CT. (c and d) ACC. (e and f) AAG. (g and h ACG. (i and j) ACT. (k and l) ATC.

After hybridization, simultaneous and sequential FISH was performed on the same cells with the repetitive probes pAs120a or pAm1, in combination with a ribosomal probe (45S or 5S) (S4-S6 Figs). This allowed all the chromosomes to be assigned to the A, C or D genomes. The nomenclature used for the reference hexaploid karyotype was that proposed by Sanz et al. [18] who used the same repetitive probes as used here. The present study confirms the locations of the ribosomal probes and the four intergenomic translocations described by the latter authors (Fig 4). Some variation was seen among metaphase cells in the detection of all chromosome pairs with C/D translocations. This also happened with the CCDD species and is related to the exposure time of the CCD camera. Certainly, the minor C/D translocation on chromosome 20D was not always evident.

The SSR-FISH results of interest with respect to the hexaploids are summarised (Fig 5) as follows: 1) The location of the hybridization signals was mostly centromeric and pericentromeric, similar to that seen found for the tetraploid species. 2) Each oligonucleotide produced signals on a few chromosomes. 3) The A genome was the best represented in the hybridization patterns, with five A chromosomes identified (8A, 11A, 13A, 15A and 16A), followed by three C genome chromosomes (2C, 4C and 6C), and three D genome chromosomes (9D, 10D and 12D). 4) As observed in the SSR hybridization patterns for the tetraploids, different oligonucleotides generated co-localized signals on several chromosomes. For example, chromosomes 10D and 15A showed a pericentromeric signal with oligonucleotides (AAC)_5_, (ATC)_5_ and (ACG)_5_, as did chromosome 2C with (AAC)_5_ and (ACT)_5_. 5) Similar to that described for tetraploid species, the SSRs located on the A and D genome chromosomes (all of them except for AAC) were not detected on the C genome chromosomes, and *vice versa*. This indicates the C genome chromosomes and the closest related A and D genomes to have different pericentromeric chromatin compositions. 6) The SSR hybridization patterns were identical for the two hexaploid species, except for CT (Figs 4a, 4b and 5) which was detected on three chromosome pairs in *A. byzantina* (8A, 13A and 9D) and on only one pair in *A. sativa* (9D). Moreover, AAC polymorphism was also observed for chromosome 6C (Figs 4c, 4d and 5).

**Fig 5.**
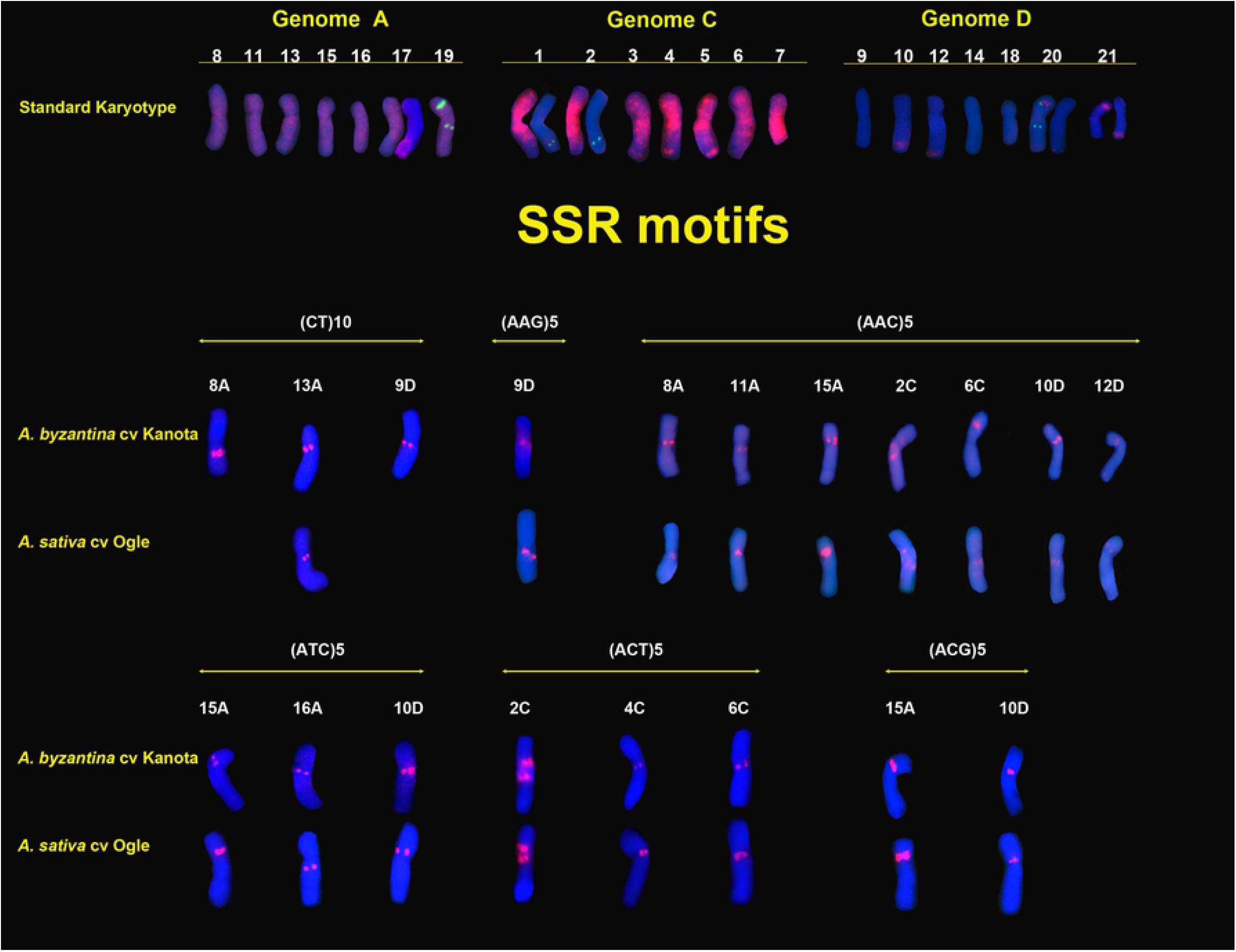
Karyotypes of hexaploid species showing a single chromosome of each homologous pair from the metaphases of cells in Fig 4. Standard karyotype is based on the FISH distribution patterns of Am1, 120a, ITS and 5S. Chromosome nomenclature is based on that Sanz et al. [18]. Standard karyotype: Am1(red), 45S (green, 19A, 20A, 21D) and 5S (green, 1C, 2C, 19D, 20D). Panels of SSR motifs show chromosomes with specific SSR signals in red.

## Discussion

### Common organization of SSR hybridization patterns in tetraploid and hexaploid species

Regions enriched in long stretches of SSR are commonly detectable by FISH. The present results indicate that most of the investigated tri-nucleotide SSR repeats do not form large blocks on the chromosomes of the three CCDD species and the two hexaploid species analyzed, in general agreement with that previously reported [39, 55–58]. Eight out of the 11 possible unique tri-nucleotide combinations were tested in this work (Table S1), and five of them (AAC, AAG, ACG, ACT and ATC) returned hybridization signals on chromosomes belonging specifically to one genome (Figs 3 and 5). This genomic preference has been observed previously for these and other SSRs in *Avena* species, although in these other reports some of the SSRs showed hybridization signals on more chromosomes than seen in the present work, and with chromosomes belonging to different genomes. For instance, Luo et al. [56, 57] described D genome chromosomes with hybridization signals for ACT and several C genome chromosomes with signals for AAC. Minor differences in the hybridization pattern for AAC were seen in the present work with respect to that reported [39, 58]. These differences might be attributable to the different methods used for inducing metaphase chromosome contraction. A greater degree of metaphase condensation was achieved by the method used in the all these authors’ work, which might have concentrated small, physically distant SSR stretches; in less condensed chromatin these stretches might not have been revealed. All these results, however, underscore differences between the C and D genome chromosomes consistent with the known strong divergence between these genomes (22, 26, 33, 34, 66].These findings are confirmed by the results of a survey of repetitive sequences extracted from *Avena*, which found some were common to the three genomes present in hexaploids, while others were specific to individual genomes or shared by the A and D genomes. However, none were shared by the C and A, or C and D genomes [66].

The tri-nucleotide SSRs investigated, and the di-nucleotide CT, located preferentially to the centromeric and pericentromeric regions in the five species studied (Figs 3 and 5). However, different SSRs or different combinations of SSRs were observed on each individual chromosome. Various SSRs co-localized on chromosomes within a species, but not always with similar intensity. For instance, AAC and ACG on chromosomes SM5 of *A. insularis*, 11D of *A. maroccana* and 10D of the hexaploids, were present at the same locations with similar intensities, suggesting they may be evenly distributed in an intermingled manner. However, these two same SSRs showed variations in intensity on different chromosome. e.g., on chromosome 7D of *A. murphyi* and 6D of *A. maroccana*, suggesting that the genomic distribution of these SSRs would be one of proximity than intermixing. Each motif would then be amplified independent of the other.

Information from the *Avena* sativa OT3098 genome assembly v1, PepsiCo (2020) https://wheat.pw.usda.gov/GG3/graingenes_downloads/oat-ot3098-pepsico) confirms that both kinds of organization are present in different regions of the chromosomes of *A. sativa*. The predicted integration map based on that assembly shows a putative pericentromeric region in chromosome 3C with two well separated clusters, each containing different SSRs [58]. Each separate cluster might be amplified independently. In contrast, mixed SSRs in a cluster would be amplified jointly. The present results suggest that an SSR combination need not necessarily be arranged in the same way on all chromosomes of an *Avena* species, as suggested by other authors [47, 51, 39]. Interestingly, the clusters integrating repeats of different SSRs, for example, AAC, ATC and ACG, were clearly maintained in specific chromosomes of the tetraploid species and hexaploids, suggesting the existence of a close relationship among these chromosomes (Figs 3 and 5).

The molecular nature of the *Avena* pericentromeric chromatin remains elusive; systematic DNA sequencing of these genomic regions has not yet been performed. In many higher eukaryotic organisms, satellite DNA is often abundant in centromeric regions, whereas the surrounding areas of pericentric chromatin more frequently contain transposon elements (TE) (mainly retrotransposon sequences) [61, 62]. Information derived from individual barley BAC clones containing SSRs indicates these sequences to be adjacent and intermingled with retrotransposon-derived sequences [63]. Given the evidence for TEs in the origin of other repetitive sequences, such as centromeric satellites [64 65], it cannot be ruled out that some SSRs originated from TE sequences. Moreover, the relationship between TEs and SSRs offers an explanation as to the distribution of SSRs in different pericentromeric regions: any SSR might be mobilized together with an adjacent TE, as proposed for satellites [62]. Further, TEs might be also involved in the distribution of SSRs, such as AC repeats, localized in other chromosome regions far from centromeres. In *Avena*, numerous families of dispersed repetitive DNA sequences localized throughout the genome have been described [66]. Retroelements are the major component of the genome of *A. sativa* genome, with Ty3/Gypsy elements representing more than 40% of all the DNA, and Ty1/Copia elements representing 5%. The genomic instability generated by allopolyploidy likely promotes TE activity, resulting in the movement of these elements along with any associated SSRs from one chromosome to another, followed by the proliferation of these SSRs in each location [67].

The poor hybridization patterns observed in *Avena* for the SSRs studied in the present work are clearly different to those described for other Poaceae species. Together with other genome information, this reveals features of *Avena* chromosome organization. For example, oligonucleotides containing motifs such as AAG, ACG or ATC in *Triticum* and *Hordeum* produce strong hybridization signals on the chromosomes of their different genomes, in agreement with estimates of the SSR frequencies (51, 52, 53, 68]. However, the high SSR frequencies in hexaploid oat genomes [42, 44] are in clear disagreement with the paucity of SSR-FISH signals described here. In the latter authors’ work, a large number of clones from enriched libraries for motifs such as ACT and ATA were obtained, but these SSRs do not seem to form large blocks of repeats as neither FISH signals were detected with (ATA)_5_ in the present work, and fewer than expected were detected with (ACT)_5_. The present results agree with those of Yan et al. [39] who also observed little abundance of A/T rich SSRs in CCDD species. The *Avena* species also lacked SSR clusters detectable by FISH that contained C/G motifs, such as CCT or CGG. These SSRs showed a highly dispersed hybridization pattern on all chromosomes of *A. insularis* and *A. sativa* with no discrete signals (Table S1; data not shown). Taken together, these results suggest that the slippage mechanism proposed to explain the amplification of repeated sequences at individual locations [69] is at work in *Avena* less intensely than in other members of Poaceae.

### Phylogenetic relationships among polyploids

The close relationships among the CCDD *Avena* species is revealed by their similarities in the chromosome distribution of the SSRs and ribosomal repeats, and their gross chromosome translocations (Figs 3, S1 Fig). This is in agreement with FISH results for 45S and 5S and Am1 [15, 26] and with those of GISH when using *A. eriantha* DNA as a probe [22, 25]. As previously discussed, clusters formed by most SSRs in any CCDD tetraploid species were present in the chromosomes of the other species, suggesting that they already were in the ancestral genome of the extant CCDD species. These results do not support the hypothesis that various allotetraploid events took place involving the participation of different diploid species as donors of the D genome - unlike that suggested by phylogenetic studies analyzing sequences of specific nuclear genes and plastic genome fragments [35]. In contrast, the present results agree with those based on genotyping by sequencing markers, which are consistent with the hypothesis that these three species likely derived from a common ancestral tetraploid [37, 38]. There are, however, interspecific differences in the chromosome structure and distribution of several repeated sequences which reveal the species-specific amplification/deletion processes that occurred during speciation. For example, the distribution of CT, AAC or ACG were very variable in the species studied. SSR distribution differences were especially significant for *A. murphyi* with respect to *A. insularis* and *A. maroccana* (Fig 3). The larger number of AAC clusters in *A. murphyi* compared to the other two species, and the lack of AC and AAG clusters in *A. murphyi*, suggest that the D genome of *A. murphyi* has been subjected to amplification processes not shared by the common ancestor of the other two species. Other studies with different SSRs also show little similarity among the SSR hybridization signals of *A. murphyi* and those of the other tetraploids [39, 54]. These observations are in agreement with the major differences in chromosome morphology and C-banding patterns observed in the karyotypes of *A. murphyi*, *A. insularis* and *A. maroccana* [5, 9, 10]. The different location of one 5S locus in *A. murphyi* compared to the other species is also noteworthy (Fig 3). The 5S locus on chromosome 7D is close to a translocated segment from a C chromosome. The likely homologous chromosomes of the other two CCDD species share the C/D translocation but not the 5S locus, suggesting that in the ancestor of these three species the translocated segment from the C genome chromosome did not encompass the entire cluster of 5S repeats. During the separate evolution of *A. murphyi*, the independent amplification of 5S sequences probably then occurred in the two chromosomes involved in the C/D translocation. 5S repeat amplification/deletion events are widely documented. A molecular diversity study of the 5S rDNA found that *A. murphyi* enclosed a 5S sequence related to genome D of *A. sativa* [70]. This was not present in *A. maroccana*, suggesting a molecular divergence in the composition of 5S repeats between the two CCDD species. Interestingly, the C/D intergenomic translocations of hexaploid species and the chromosome locations of 5S loci in these species are the same as those in *A. insularis* and *A. maroccana* (Figs 3 and 5). Other studies have also shown differences in the number of minor 45S rDNA loci in *A. murphyi* with respect to the other two CCDD species [10]. Together, these cytogenetic data indicate that *A. murphyi* may have undergone early chromosome differentiation from the other two CCDD species, ruling it out as being directly involved in the origin of the hexaploids.

As pointed out by Yan et al. [39] and as mentioned in the introduction to this paper, no agreement has been reached regarding which tetraploid species may have contributed to the hexaploid oat genome. From a cytogenetic point of view, identifying karyotypic similarities possibly indicative of homology would help understanding the relationships among phylogenetically close species. In this endeavour, the rich pattern of signals returned by oligonucleotide (AC)_10_ in D genome chromosomes of has been of great use (Fig 1). For purposes of comparison among karyotypes, a representation of the hybridization signals of *A. insularis, A. maroccana, A. sativa/A. byzantine* is shown (Fig 6). The information on the AC hybridization profiles from *A. maroccana* and the hexaploids is taken from Fominaya et al. [54].

**Fig 6.**
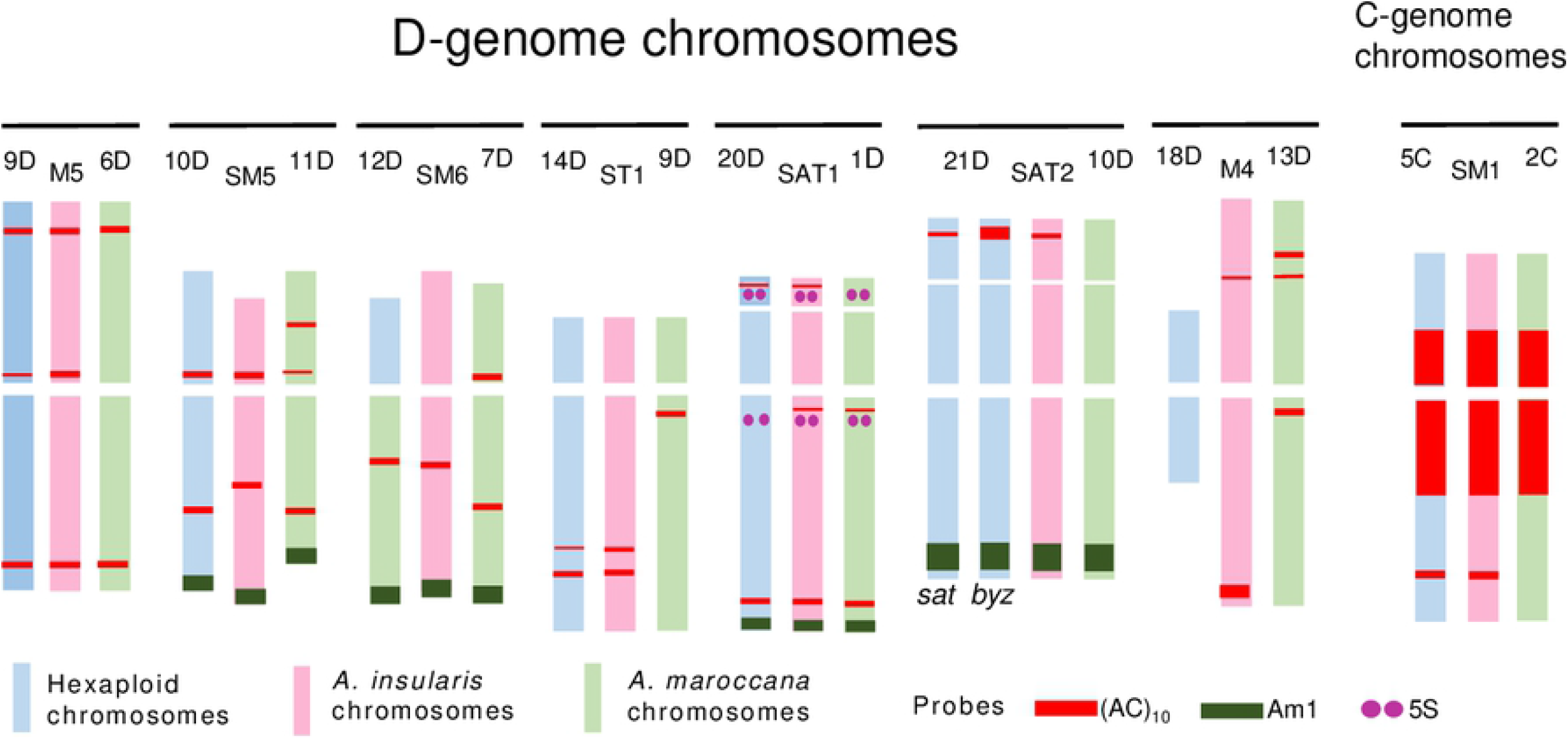
Schematic drawing of oligonucleotide (AC)_10_ FISH patterns (red) showing all chromosomes of genome D and chromosome 5C of hexaploids, (*A.sativa* and *A. byzantina*), and corresponding chromosomes of *A. insularis* and *A. maroccana*. Loci for 5S (purple) and signals for Am1 (green) on translocated C/D chromosomes are also represented. FISH patterns of hexaploids and *A. maroccana* were taken from A. Fominaya et al. [54]. Karyotypes and chromosome nomenclatures were those of Sanz et al. [18] for hexaploids; Fominaya et al. [5, 54] for *A. marocanna* and Jellen and Ladizinsky [9] for *A. insularis*. (AC)_10_ polymorphism between *A. sativa* and *A. byzantina* for chromosome 21D is indicated.

Great conservation of the interstitial AC signals was seen in the D genome chromosomes of the different species, and of the other SSRs studied here in the pericentromeric areas of the C and D genome chromosomes (Figs 3 and 6). The polymorphism detected for several SSR hybridization signals between *A. insularis* and *A. maroccana* - both in terms of the presence/absence and signal intensity - was very useful for discerning which of these two extant species was more closely related to the hexaploid taxa, and the present results suggests the likely involvement of *A. insularis* in the origin of the hexaploids (Figs 3, 5 and 6). Certainly, not all the SSR hybridization signals present in *A. insularis* were exactly coincident with those observed in the putative homologous of hexaploids, but the SSR hybridization patterns for *A. insularis* matched those observed for the hexaploid species much better than did those of *A. maroccana*. Indeed, the hybridization patterns of AC, CT and ACG of *A. insularis* and *A. maroccana* were quite different in terms of the number of chromosomes and/or the specific chromosomes bearing the signals, whereas they were mostly coincident in *A. insularis* and the hexaploids. Moreover, the AAC and ATC signals were more intense in *A. maroccana* than in *A. insularis* (on the related chromosomes M5 and 6D, respectively). These signals were, however, missing in the hexaploids, indicating deletions of these SSRs in the lineage from *A. insularis* to the hexaploids. The hybridization patterns of CT and AC in *A. insularis* in particular suggest a close relationship of this species with *A. byzantina* (Figs 3, 5 and 6). Interestingly, prominent AC hybridization signals were only observed on the long arm of chromosome M4 of *A. insularis*, and on the short arm of 21D of *A. byzantina*. None of the other analyzed species returned a similar signal (Figs 1 and 6). In addition, none of the tetraploid taxa showed a chromosome as small and metacentric as chromosome 18D of the hexaploids (Fig 3 and 5), indicating this chromosome came from an ancestral form that suffered an important reduction in size, which likely occurred after the alloploidyzation leading to the hexaploid progenitor. Thus, all the hexaploid taxa 18D chromosomes share this morphology [13]. This size reduction might be partly explained by a translocation from the long arm of chromosome 4M of *A. insularis* encompassing the large terminal AC cluster, to the short arm of chromosome 21D of the hexaploid progenitor. In both chromosomes the size of the AC signal is very similar. Thereafter, *A. byzantina* would have maintained the AC block on 21D, whereas in *A. sativa* AC sequences would have been deleted or rearranged in an independent event. To explain the differences between *A. byzantina* and *A. sativa* with respect to chromosome 21D, a chromosome reorganization between chromosomes 21D and 4C was postulated [54], since 4C of *A. sativa* also has a terminal AC signal, although smaller than that of 21D of *A. byzantina*. This indicates a partial deletion of the AC cluster in chromosome 4C of *A. sativa*. Interestingly, chromosome 21D participates in a translocation common to all the hexaploid species [18] and in another cultivar-specific translocation involving chromosome 17A [54]. The important role played by chromosome translocations in the evolution of the *Avena* genus is widely documented. Moreover, the isolation of new repeated sequences and their location by FISH have led to several new minor intra- and intergenomic translocations being found in the hexaploid genomes with respect to the diploid taxa [58, 66]. However, the use of different and poorly correlated chromosome nomenclatures between the latter articles and the present prevent the carrying out of tests to determine whether chromosome 21D is prone to suffer such rearrangements.

According to Loskutov [71] and Ladizinsky [21] who reviewed the origin of hexaploid oats based on geographical distribution, botanical features and the chromosome pairing of hybrids, the progenitor of all hexaploids is *A. sterilis* - or a closely related form. From it, two branches evolved separately. One led to *A. byzantina* and the second to *A. occidentalis* and the other hexaploids, including *A. fatua* and *A. sativa*. This early separation of *A. byzantina* from *A. sativa* may have been accompanied by structural genome rearrangements - not all of them well identified cytologically but detected by the presence of several distorted linkage groups in the genetic map derived from crosses between these two hexaploids [72]. If the above hypothesis holds, it is likely that the primitive form of *A. sterilis* carried an AC signal on chromosome 21D. During the present work, one *A. sterilis* accession (data not shown) was studied but it failed to show this AC hybridization signal. However, Badaeva et al. [13] observed polymorphism for the C-banding patterns of *A. sterilis*, especially on the short arm of this chromosome. The study of this and other accessions might help determine whether this C banding polymorphism matches the AC hybridization signal observed in *A. insularis* and *A. byzantina*. If so, the present results suggesting *A. insularis* to be the closest ancestor of hexaploids would be cytogenetically reinforced, strongly supporting those of genomic studies on the role of this species in the origin of hexaploid oats.

Taken together our cytogenetic results on *A. insularis* and *A. maroccana*, it is feasible to relate each D-genome chromosome of one species with its putative homologous of the other, also with the corresponding hexaploid chromosome. Although less chromosome markers were obtained for C-genome chromosomes, presumed relationships can be reached for several of these chromosomes. Based on these data, a common nomenclature for chromosomes of these species is proposed (Table 2). This should help to future works that could fine tune better the extension of homology among chromosomes of these species.

**Table 2.** Common nomenclature for *A. insularis* and *A. maroccana* for D-genome chromosomes and several C-genome chromosomes regarding to chromosomes of the hexaploid species, based on SSR hybridization patterns shared for at least two species, 45S and 5S loci, and intergenomic translocations (C/D or D/C).

| *A. insularis* | *A. maroccana* | Common nomenclature | SSR | rDNA | C/D or D/C |
| --- | --- | --- | --- | --- | --- |
| M5 | D6 | 9D | AC, AAC, ATC, CT | - | - |
| SM5 | 11D | 10D | AC, AAC, ACG | - | C/D |
| SM6 | 7D | 12D | AC | - | C/D |
| ST1 | 9D | 14D | AC | - | - |
| M4 | 13D | 18D | AC, AAG | - | - |
| SAT1 | 1D | 20D | AC | 45S, 5S | C/D |
| SAT2 | 10D | 21D | AC | 45S | C/D |
| M1 | 3C | 1C | - | 5S | D/C |
| M2 | 5C | 2C | ACT | 5S | D/C |
| SM2 | 4C | 4C | ACT | - | - |
| SM1 | 2C | 5C | AC | - | - |
| SM3 | 12C | 6C | ACT | - | - |

## Conclusions

FISH hybridization patterns of seven SSRs, two ribosomal repeated sequences, and a C genome-specific repetitive DNA sequence, showed common hybridization signals on chromosomes with similar morphologies in three CCDD and two AACCDD species of *Avena*. The similarities among chromosomes of different species allowed tentative homologous relationships to be established among all D genome chromosomes and several C genome chromosomes of *A. insularis, A. maroccana, A. sativa* and *A. byzantina*, highlighting the close genetic relationships among them. In contrast, *A. murphyi* showed obvious differences in its SSR hybridization signals, and in its entire karyotype, suggesting it evolved somewhat separately from the other two CCDD species. The few but significant differences in the SSR hybridization patterns of *A. insularis* and *A. maroccana* helped to establish that *A. insularis* is more likely to be involved in the origin of the hexaploids than is *A. maroccana*. *A. insularis* shared diagnostic FISH signals with the hexaploids, especially with *A. byzantina*. The present results support the hypothesis that the extant *A. insularis*, or its direct ancestor, is a strong candidate as the progenitor of hexaploid oats.

## Supporting information

**S1 Table**. Occurrence and main distribution of SSRs in CCDD tetraploids: *A. insularis, A. maroccana* and *A. murphyi* and AACCDD hexaploids: *A. byzantina* and *A. sativa*.

**S1 Fig**. FISH of mitotic metaphases of tetraploid species showing C/D intergenomic translocations (arrows).

**S2 Fig.** FISH of mitotic metaphases of CCDD tetraploid species showing the distribution of SSRs CT, AAG and AAC. When positive signals for SSRs were observed, the same cells rehybridized with pAm1 are shown. (a and b) *A. insularis*. (c) *A. maroccana*. (d) *A. murphyi*. (e and f) *A. insularis*. (g and h) *A. maroccana*. (i) *A. murphyi*. (j and k) *A. insularis*. (l and m) *A. maroccana*. (n and o) *A. murphyi*

**S3 Fig.** FISH of mitotic metaphases of CCDD tetraploid species showing the distribution of SSRs ATC, ACT and ACG. Same cells after rehybridization showing signals for Am1, 45S and 5S as indicated on the microphotographs. (a and b) *A. insularis*. (c and d) *A. maroccana*. (e and f) *A. murphyi*. (g and h) *A. insularis*. (i and j) *A. maroccana*. (k and l) *A. murphyi*. (m and n) *A. insularis*. (o and p) *A. maroccana*. (q and r) *A*. *murphyi*.

**S4 Fig.** FISH of mitotic metaphases of hexaploid species showing the distribution of SSRs ACG and ACT. Same cells after rehybridization showing signals for Am1, 120a, 45S and 5S as indicated on the microphotographs. (a-c) *A. byzantina*. (d-f) *A. sativa*. (g-i) *A. byzantina*. (j-l) *A. sativa*.

**S5 Fig.** FISH of mitotic metaphases of hexaploid species showing the distribution of SSRs CT and AAG. Same cells after rehybridization showing signals for Am1, 120a, 45S and 5S as indicated on the microphotographs. (a-c) *A. byzantina*. (d-f) *A. sativa*. (g-i) *A. byzantina*. (j-l) *A. sativa*.

**S6 Fig.** FISH of mitotic metaphases of hexaploid species showing the distribution of SSRs AAC and ATC. Same cells after rehybridization showing signals for Am1, 120a, 45S and 5S as indicated on the microphotographs. (a-c) *A. byzantina*. (d-f) *A. sativa*. g-i) *A. byzantina*. (j-l) *A. sativa*.

## Acknowledgments

We thank University of Alcalá (IUICP/PI2019/001) for supporting this work. We also thank to Juan M. Barrero for technical assistance.

## Notes

**Competing interest**, The authors have declared that no competing interests exist.

### Competing Interest Statement

The authors have declared no competing interest.

## References

1. Thomas H. Cytogenetics of *Avena*. Oat science and technology. Agronomy, Monograph No 33. Edited by HG Marshall and ME Sorrells. ASA, CSSA, SSA, Madison, Wisc. pp. 473–507. 1992.

2. Loskutov I, Rines HW. Avena. Wild crop relatives: genomic and breeding resources: cereals. Springer, New York. Pp 109–184. 2011.

3. Rajhathy T, Thomas H. Genetic control of chromosome pairing in hexaploid oats. Nat New Biol. 1972; 239:217–219. https://doi.org/10.1038/newbio239217a0

4. Fominaya A, Vega C, Ferrer E. Giemsa C-banded kayotypes of *Avena* species. Genome. 1988; 30:627–631. https://doi.org/10.1139/g88-106

5. Fominaya A, Vega C, Ferrer E. C-banding and nucleolar activity of tetraploid *Avena* species. Genome. 1988; 30:633–638. https://doi.org/10.1139/g88-107

6. Linares C, Vega C, Ferrer E, Fominaya A. Identification of C-banded chromosomes in meiosis and the analysis of nucleolar activity in *Avena byzantina* C. Koch cv Kanota. Theor Appl Genet. 1992; 83:650–654. https://doi.org/10.1007/BF00226911

7. Jellen EN, Phillips RL, Rines HW. C-banding karyotypes and polymorphisms in hexaploid oat accessions *(Avena* ssp.) using Wright’s stain. Genome. 1993; 36:1129–1137. https://doi.org/10.1130/g93-151

8. Jellen EN, Rooney WL, Phillips RL Rines HW. Characterization of hexaploid oat *Avena byzantina* cv. Kanota monosomic series using C-banding and RFLPs. Genome. 1993; 36:062–970. https://doi.org//10.1139/g93-126

9. Jellen EN, Ladizinsky G. Giemsa C-banding in *Avena insularis* Ladizinsky. Genet Resour Crop Evol. 2000; 47:227–230. https://doi.org/10.1023/A:1008769105071

10. Shelukhina OY, Badaeva ED, Loskutov IG, Pukhalsky VA. A comparative cytogenetic study of the tetraploid oat species with the A and C genomes: *Avena insularis, A. magna*, and *A*. *murphyi*. Russ J Genet. 2007; 43:747–761. https://doi.org/10.1134/S102279540706004X

11. Shelukhina OY, Badaeva EK, Loskutok IG. Comparative analysis of diploid species of *Avena* L. using cytogenetic and biochemical markers: *Avena canariensis* Baum et Fedak and *A. longiglumis* Dur. Russ J Genet. 2008; 44:694–701. https://doi.org/0.1134/S1022795408060094

12. Shelukhina OY, Badaeva ED, Brezhneva TA, Loskutov IG, Pukhalsky VA. Comparative analysis of diploid species of *Avena* L. using cytogenetic and biochemical markers: *Avena Pilosa* MB and *A. clauda* Dur. Russ. Russ J Genet. 2008; 44:1087–1091. https://doi.org/10.1134/S1022795408090111

13. Badaeva ED, Shelukhina OY Dekova OS, Loskutov IG, Pukhalsky VA. Comparative cytogenetic analysis of hexaploid *Avena* L. species. Russ J Genet. 2011; 47:691–702. https://doi.org/1134/S1022795411060068

14. Fominaya A, Hueros G, Loarce Y, Ferrer E. Chromosomal distribution of a repeated DNA sequence from C-genome heterochromatin and identification of a new ribosomal DNA locus in the *Avena* genus. Genome. 1995; 38:548–557. https://doi.org/10.1139/g95-071

15. Linares C, González JM, Ferrer E, Fominaya A. The use of double fluorescence in situ hybridization to physically map the positions of 5S rDNA genes in relation to the chromosomal locations of 18S-5.8S-26S rDNA and a C genome specific sequence in the genus *Avena*. Genome. 1996; 39:535–542. https://doi.org/10.1139/g96-068.

16. Linares C, Ferrer E, Fominaya A. Discrimination of the closely related A and D genomes of the hexaploid oat *Avena sativa* L. Proc Natl Acad Sci. 1998; 95: 12450–12455. https://doi.org/10.1073/pnas.95.21.12450

17. Irigoyen ML, Loarce Y, Linares C, Ferrer E, Leggett JM, Fominaya A. Discrimination of the closely relate A and B genomes in tetraploid species of *Avena*. Theor Appl Genet. 2001; 103:1160–1166. https://doi.org/10.1007/s001220100723

18. Sanz MJ, Jellen EN, Irigoyen ML, Ferrer E, Fominaya A. A new chromosome nomenclature system for oat (*Avena sativa* L. and *A*. *byzantina* C. Koch) based on FISH analysis of monosomic lines. Theor Appl Genet. 2010; 12:1541–1552. https://doi.org/10.1007/s00122-010-1409-3

19. Luo X, Xhang H, Kang H, Fan X, Wang Y, Sha L et al. Exploring the origin of the D genome of oat by fluorescence in situ hybridization. Genome. 2014; 57:469–472. https://doi.org/10.1139/gen-2014-0048

20. Luo X, Tinker NA, Zhang H, Wight CP, Kang H, Fan X et al. Centromeric position and genomic allocation of a repetitive sequence isolated from chromosome 18D of hexaploid oat *Avena sativa* L. Genet Resour Crop Evol. 2015; 62:1–4. https://doi.org/10.1007/s10722-014-0170-x

21. Ladizinsky G. Studies in oat evolution: a man’s life with Avena. Springer. Heidelberg, Germany. 2012.

22. Jellen EN, Gill BS, Cox TS. Genomic in situ hybridization differentiates between A/D and C-genome chromatin and detects intergenenomic translocations in polyploid oat species (genus *Avena*). Genome. 1994; 37:613–618. https://doi.org10.1139/g94-087

23. Chen Q, Armstrong K. Genomic in situ hybridization in *Avena sativa*. Genome. 1994; 37:607–612. https://doi.org/10.1139/g94-086

24. Leggett JM, Markland GS. The genomic structure of *Avena* revealed by GISH. Kew Chromosome Conference IV, London. Edited by PE Brandham and MD Bennett, pp. 133–139. 1995.

25. Hayasaki M, Morikawa T, Tarumoto I. Intergenomic translocations of polyploid oats (genus *Avena)* revealed by genomic in situ hybridization. Genes Genet Syst. 2000; 75:167–171. https://doi.org/10.1266/ggs.75.167.

26. Linares C, Irigoyen ML, Fominaya A. Identification of C-genome chromosomes involved in intergenomic translocations in *Avena sativa* L. using cloned repetitive DNA sequences. Theor Appl Genet. 2000; 100:353–360. https://doi.org/10.1007/s001220050046

27. Oliver RE, Jellen EN, Ladizinski G, Korol AB, Kilian A, Beard JL et al. New diversity array technology (DArT) markers for tetraploid oat (*Avena nagna* Murphy et Terrel) provide the first complete oat linkage map and markers linked to domestication genes from hexaploidy *A. sativa* L. Theor Apll Genet. 2011; 123:1159–1171. https://doi.org/10.1007/s00122-011-1656-y.

28. Chaffin AS, Huang YF, Smith S, Bekele WA, Babiker E, Gnaesh BN et al. A consensus map in cultivated oat revealed conserved grass synteny with substantial sub-genome rearrangement. Plant Genome 2016; 9:2. https://doi:10.3835/plantgenome2015.10.0102

29. Latta RG, Bekele WA, Wight CP, Tinker NA. Comparative linkage mapping of diploid, tetraploid, and hexaploid *Avena* species suggests extensive chromosome rearrangement in ancestral diploids. Sci Rep. 2019; 9:12298. https://doi,org/10.1038/s41598-019-48639-7

30. Katsiotis A, Schmidt T, Heslop-Harrison JS. Chromosomal and genomic organization of *Ty-copia-like* retrotransposon sequence in the genus *Avena*. Genome. 1995; 39:410–417. https://doi.org/10.1139/g96-052

31. Ladizinsky. A new species of *Avena* from Sicily, possibly the tetraploid progenitor of hexaploid oats. Genet Resour Crop Evol. 1998; 45:263–269. https://doi.org/10.1023/A:1008657530466

32. Ladizinsky G, Zohary D. Notes on species delimitation, species relationships and polyploidy in *Avena* L. Euphytica. 1971; 20:380–395. https://doi.org/10.1007/BF00035663

33. Fominaya A, Loarce Y, Irigoyen ML, Ferrer E. Molecular genetic and cytogenetic evidences supporting the genomic relationships of the genus *Avena.* Plant genome, biodiversity and evolution. Vol 1 part D. Phanerogams (Gymnosperm) ad (Angiosperm-Monocotyledons). Edited by: AK Sharma, A. Sharma. Sci Pub. USA. Pp 305–321. ISBN 978-157808-402-3. 2006.

34. Yan H, Baum BR, Zhou P, Wei Y, Ren C, Xiong F, et al. Phylogenetic analysis of the genus *Avena* based on chloroplast intergenic spacer *psb*A–*trn*H and single-copy nuclear gene Acc1. Genome. 2014; 57:267–277. https://doi.org/10.1139/gen-2014-0075

35. Liu Q, Lin L, Zhou X, Peterson P, Wen J. Unraveling the evolutionary dynamics of ancient and recent polyploidization events in *Avena* (Poaceae). Sci Rep. 2017; 7: 41944. https://doi.org/10.1038/srep41944

36. Fu Y. Oat evolution revealed in the maternal lineages of 25 *Avena* species. Sci Rep. 2018; 8:4245. https://doi.org/10.1038/s41598-018-20586-9

37. Chew P, Meade K, Hayes A, Harjes C, Bao Y, Beattie AD, et al. A study on the genetic relationships of *Avena* taxa and the origins of hexaploid oat. Theor Appl Genet. 2016; 129:1405–1415. https://doi.org/10.1007/s00122-016-2712-4

38. Yan H, Bekele W, Wight C, Peng Y, Langdon T, Latta R, et al. High-density marker profiling confirms ancestral genomes of *Avena* species and identifies D-genome chromosomes of hexaploid oat. Theor Appl Genet. 2016; 129: 2133–2149. https://doi.org/10.1007/s00122-016-2762-7

39. Yan H, Ren Z, Deng D, Yang K, Yang C, Zhou P, et al. New evidence confirming the CD genomic constitutions of the tetraploid *Avena* species in the section *Pachycarpa* Baum. PLoS ONE. 2021; 16(1): e0240703. https://doi.org/10.1371/journal.pone.0240703

40. Peng Y, Wei Y, Baum BR, Yan Z, Lan X, Dai S, et al. Phylogenetic inferences in *Avena* based on analysis of *FL* intron2 sequences. Theor App Genet 2010; 121:985–1000. https://doi.org/10.1007/s00122-010-1367-9

41. Peng Y, Zhou P, Zhao J, Li J, Lai S, Tinker NA, et al. Phylogenetic relationships in the genus *Avena* based on the nuclear Pgk1 gene. PLoS ONE. 2018; 13(11): e0200047. https://doi.org/10.1371/journal.pone.0200047

42. Li CD, Rossnagel BG Scoles GJ. The development of oat microsatellite markers and their use in identifying relationship among *Avena* species and oat cultivars. Theor Appl Genet. 2000;1259–1268. https://doi.org/10.1007/s001220051605

43. Becher R. EST-derived microsatellites as a rich source of molecular markers for oats. Plant Breeding 2007; 126:274–278. https://doi.org/10.111/j.1439-0523.2007.01330.x

44. Song G, Huo P, Wu B, Zhang Z. A genetic linkage map of hexaploid naked oat constructed with SSR markers. Crop J. 2015;353–357. https://doi.org/10.1016/j.cj.2015.01.005

45. Dumlupinar Z, Brown R, Campbell R, Jellen EN, Anderson J, Bonman JM et al. The art of attrition: development of robust oat microsatellites. Plant Breeding 2016; 135:323–334. htpps://doi.org/10.1111/pbr.12362

46. Cuadrado A, Schwarzacher T. The chromosomal organization of simple sequence repeats in wheat and rye genomes. Chromosoma. 1998; 107(8):587–594. https://doi.org/10.1007/s004120050345

47. Cuadrado A, Cardoso M, Jouve N. Physical organization of simple sequence repeats (SSRs) in Triticeae: structural, functional and evolutionary implications. Cytogenet Genome Res. 2008; 120:210–219. https://doi.org/10.1159/000121069.

48. Zheng J, Sun C, S Z, Hou X, Bonnema G. Cytogenetic diversity of simple sequences repeats in morphotypes of *Brassica rapa* ssp. *chinensis*. Front Plant Sci. 2016; 7:1049. https://doi.org/10.3389/fpls.2016.01049

49. Elder JF, Turner BJ. Concerted evolution of repetitive DNA sequence in eukaryotes. Q Rev Biol. 1995; 70:297–320. https://doi.org/10.1086/419073

50. López-Flores I, Garrido-Ramos MA. The repetitive DNA content of eukaryotic genomes. Genome Dyn. 2012; 7:1–28. htpps://doi.org/10.1159/000337118

51. Zhang, Y., Fan, C., Chen, Y. et al. Genome evolution during bread wheat formation unveiled by the distribution dynamics of SSR sequences on chromosomes using FISH. BMC Genomics 2021; 22, 55. https://doi.org/10.1186/s12864-020-07364-6

52. Carmona A, Friero E, de Bustos A, Jouve N, Cuadrado A. Cytogenetic diversity of SSR motifs within and between *Hordeum* species carrying the H genome: *H*. *vulgare* L. and *H*. *bulbosum* L. Theor Appl Genet. 2013; 126:949–961. https://doi.org/10.1007/s00122-012-2028-y

53. Dou Q, Liu R, Yu F. Chromosomal organization of repetitive DNAs in *Hordeum bogdanii* and *H. brevisubulatum* (Poaceae). Comp Cytogenet. 2016; 7;10:465–481. https://doi.org/10.3897/CompCytogen.v10i4.9666.

54. Fominaya A, Loarce Y, Montes A, Ferrer E. Chromosomal distribution patterns of the (AC)10 microsatellite and other repetitive sequences, and their use in chromosome rearrangement analysis of species of the genus *Avena*. Genome. 2017; 60:216–227. https://doi.org/10.1139/gen-2016-0146

55. Luo X, Tinker NA, Zhou Y, Liu J, Wan W, Chen LJAPP. A comparative cytogenetic study of 17 *Avena* species using Am1 and (GAA)6 oligonucleotide FISH probes. Acta Physiol Plant 2018; 40:145. https://doi.org/10.1007/s11738-018-2721-9

56. Luo X, Tinker NA, Zhou Y, Liu J, Wan W, Chen LJGR, et al. Chromosomal distributions of oligo-Am1 and (TTG)6 trinucleotide and their utilization in genome association analysis of sixteen *Avena* species. Genet Resour Crop Evol. 2018; 65:1625–1635.https://doi.org/10.1007/s10722-018-0639-0

57. Luo X, Tinker NA, Zhou Y, Wight CP, Liu J, Wan W, et al. Genomic relationships among sixteen *Avena* species based on (ACT)6 trinucleotide repeat FISH. Genome. 2018; 61:63–70. https://doi.org/10.1139/gen-2017-0132

58. Jiang, W., Jiang, C., Yuan, W. et al. A universal karyotypic system for hexaploid and diploid *Avena* species brings oat cytogenetics into the genomics era. BMC Plant Biol 2021; 21, 213. https://doi.org/10.1186/s12870-021-02999-3

59. Solano R, Hueros G, Fominaya A, Ferrer E. Organization of repeated sequences in species of the genus *Avena*. Theor Appl Genet. 1992; 83:602–607. https://doi.org/10.1007/BF00226904

60. Gerlach WL, Dyer TA. Sequence organization of the repeating units in the nucleus of wheat which contain 5S rRNA genes. Nucleic Acids Res. 1980; 8:4851–65. http://doi.org/10.1093/nar/8.21.4851.

61. Gong Z, Wu Y, Koblízková A, Torres GA, Wang K, Iovene M, Neumann P, Zhang W, Novák P, Buell CR, Macas J, Jiang J. Repeatless and repeat-based centromeres in potato: implications for centromere evolution. Plant Cell. 2012; 24:3559–74. http://doi.org/10.1105/tpc.112.100511.

62. Hartley G, O’Neill RJ. Centromere Repeats: Hidden Gems of the Genome. Genes (Basel). 2019; 16:10:223. https://doi.org/10.3390/genes10030223.

63. De Bustos A, Cuadrado C Jouve N. Sequencing of long stretches of repetitive DNA. Sci. Rep. 2016; 6, 36665. https://doi.org/10.1038/srep36665.

64. Smýkal P, Kalendar R, Ford R, Macas J, Griga M. Evolutionary conserved lineage of *Angela*-family retrotransposons as a genome-wide microsatellite repeat dispersal agent. Heredity 2009; 103:157–67. https://doi.org/10.1038/hdy.2009.45

65. Paço A, Freitas R, Vieira-da-Silva A. Conversion of DNA Sequences: From a Transposable Element to a Tandem Repeat or to a Gene. Genes (Basel). 2019; 5: 10:1014. https://doi.org/10.3390/genes10121014

66. Liu, Q., Li, X., Zhou, X. et al. The repetitive DNA landscape in *Avena* (Poaceae): chromosome and genome evolution defined by major repeat classes in whole-genome sequence reads. BMC Plant Bio 2019;19, 226. https://doi.org/10.1186/s12870-019-1769-z

67. Han J, Masonbrink RE, Shan W, Song F, Zhang J, Yu W, et al. Rapid proliferation and nucleolar organizer targeting centromeric retrotransposons in cotton. Plant J. 2016; 88: 992–1005. https://doi,org/10.1111/tpj.13309

68. Adonina IG, Goncharov NP, Badaeva ED, Sergeeva EM, Petrash NV, Salina EA. (GAA)n microsatellite as an indicator of the A genome reorganization during wheat evolution and domestication. Comparative Cytogenet. 2015. 9:533 47. https://doi.org/10.3897/CompCytogen.v9i4.5120

69. Schlotterer C. Evolutionary dynamics of microsatellite DNA. Chromosoma 2000; 109: 65–371. htpps://doi.org/10.1007/s004120000089.

70. Peng YY, Wei YM, Baum BR, Zheng YL. Molecular diversity of the 5S rRNA gene and genomic relationships in the genus *Avena* (Poaceae: Aveneae). Genome. 2008; 512:137–54. https://doi.org/10.1139/g07-111

71. Loskutov IG. On evolutionary pathways of *Avena* species. Genet Resour Crop Evol. 2008; 55:211–220. https://doi.org/10.1007/s10722-007-9229-2

72. Tinker NA, Kilian A, Wight CP, Heller-Uszynska K, Wenzl P, Rines HW, et al. New DArT markers for oat provide enhanced map coverage and global germplasm characterization. BMC Genomics 2009; 10, 39. https://doi.org/10.1186/1471-2164-10-39

